# LDL receptor-mediated endocytosis of *Escherichia coli* α-hemolysin mediates renal epithelial toxicity

**DOI:** 10.1101/2025.02.14.638193

**Authors:** Hunter W. Kuhn, Madeleine R. Smither, Rachel J. Jin, Christina A. Collins, Hongming Ma, Jason Sina, Joseph P. Gaut, Michael S. Diamond, David A. Hunstad

## Abstract

Pore-forming toxins (PFTs) are secreted bacterial effector molecules that disrupt host cell membranes. The α-hemolysin (HlyA) of uropathogenic *Escherichia coli* (UPEC) can exert damage to various mammalian cell types. While a candidate toxin receptor (CD11a/CD18 [LFA-1] integrin) exists on myeloid cells, the mechanism of HlyA cytotoxicity to epithelial cells remains undefined. We show that HlyA secretion by UPEC exacerbates renal tubular epithelial injury during ascending pyelonephritis in mice. A CRISPR-Cas9 loss-of-function screen in renal collecting duct cells identified clathrin-mediated endocytosis as required for HlyA cytotoxicity. HlyA internalization induces lysosomal permeabilization, facilitating protease release, cytoplasmic acidification, and mitochondrial dysfunction leading to rapid cell death. This mechanism contrasts with the described actions of other PFTs (plasma membrane poration and osmotic cytolysis). We also identify the low-density lipoprotein receptor (LDLR) as an epithelial receptor for HlyA; genetic or competitive inhibition of the HlyA-LDLR interaction prevented cytotoxicity. Our studies define a new mechanism of action for HlyA, in which its toxicity to epithelial cells requires LDLR-mediated, clathrin-dependent internalization. These results suggest therapeutic avenues for mitigating HlyA-induced damage during *E. coli* infections.

## INTRODUCTION

Bacterial toxins are central to pathogenesis and inflict host tissue damage through direct cell injury and by provoking deleterious inflammatory responses. Strains of uropathogenic *Escherichia coli* (UPEC) encode for multiple secreted toxins that contribute to host inflammation and renal damage, including α-hemolysin (HlyA), a prototypic member of the large bacterial repeats-in-toxin (RTX) family, so named for a conserved GGxGxDxUx sequence repeat.^1^ UPEC is the primary cause of urinary tract infections (UTI), both in the bladder (cystitis) or the kidney (pyelonephritis). Pyelonephritis can leave renal scars despite appropriate antibiotic treatment^2–4^ and confers risk for chronic kidney disease in later life.^5,6^ Carriage of the *hlyCABD* operon, which encodes for the active toxin (HlyA) as well as modification and delivery components, is more prevalent among UPEC isolates from patients with pyelonephritis than with cystitis^7,8^. Moreover, there is wide variation in hemolytic activity (*i.e*., HlyA secretion) among UPEC strains, reflecting complex transcriptional activation and *hlyCABD* coding polymorphisms.^9^ Though recent advances in mouse models have enabled more detailed interrogation of bacterial pathogenesis in the kidney, the influence of HlyA in the pathogenesis of pyelonephritis remains poorly defined.

HlyA is secreted via classical Gram-negative type 1 secretion^10,11^ and can exert cytotoxicity in a variety of cell types. Typical structural and biochemical characterization of HlyA has been precluded by long-standing challenges in purification and functional analysis, due largely to its amphipathic nature and poor stability.^12^ As HlyA can lyse red blood cells (RBCs),^13^ and biochemical (though not structural) data indicate it can form pores in erythrocyte membranes,^14,15^ HlyA has long been presumed to function as a pore-forming toxin (PFT) that causes cell death via disruption of osmotic homeostasis.^1,16^ Application of sublytic concentrations of HlyA to epithelial cells led to activation of serine proteases in the cytosol of intoxicated cells (posited to be a downstream effect of plasma membrane poration)^17^ and elicited production of cytokines such as IL-6, IL-8, and GM-CSF.^18,19^ HlyA has been studied mostly in myeloid cells, where (as with several other RTX toxins) direct interaction with the leukocyte β_2_-intergrin CD11a/CD18 (LFA-1) is implicated in its cytotoxicity.^20–22^ As healthy epithelial cells lack CD11a/CD18 expression, HlyA must function in ways beyond those currently known.

Leveraging an updated preclinical mouse model of ascending pyelonephritis, we show that HlyA acts to exacerbate renal tubular injury *in vivo*. Utilizing a CRISPR-Cas9 loss-of-function screen, we show that HlyA enters epithelial cells via clathrin-mediated endocytosis (CME) and exerts its cytotoxicity via a mechanism distinct from the proposed model of plasma membrane poration. Instead, HlyA triggers lysosomal permeabilization, leading to rapid cytoplasmic acidification and protease release, with ensuing mitochondrial dysfunction and caspase-independent cell death. Furthermore, we identify the low-density lipoprotein receptor (LDLR) as an HlyA receptor. Genetic disruption of *Ldlr* reduced HlyA binding to murine renal epithelial cells *in vitro* and blocked HlyA-induced cell death. Introduction of a soluble LDLR ectodomain fusion peptide (LDLR-Fc) or competition with exogenous LDL effectively neutralized HlyA toxicity against renal epithelial cells. Our experiments identify a mechanism by which HlyA interacts with and intoxicates epithelial cells, and illuminate a novel target for mitigating UPEC-associated tissue injury during UTI.

## RESULTS

### α-Hemolysin secretion enhances renal epithelial injury during ascending UTI

During ascending pyelonephritis, UPEC travel from the bladder through the ureter to access the renal collecting system and colonize the renal tubular space, provoking a local inflammatory response that contributes to renal tissue injury; this disease pathogenesis sequence has been modelled in mice.^23–27^ To determine the contribution of α-hemolysin to renal injury during pyelonephritis, we generated independent *hlyA* disruption mutants (Δ*hlyA:1,* Δ*hlyA:2*) and a chromosomally based HlyA-overexpressing strain (*hlyA^++^*) in the UPEC isolate UTI89 (**Supplementary Fig. 1**). These mutant strains or wild-type (WT) UTI89 were transurethrally inoculated into the bladders of female C57BL/6 mice (with prior exposure to testosterone, which promotes kidney infection in this background^24,25^), which were later euthanized at several time points for histological analyses and bacterial load determination (**Fig. 1A**). Consistent with prior studies,^19,28^ genetic deletion of *hlyA* had no significant impact on UPEC colonization of the kidneys (**Fig. 1B**) or bladder (**Supplementary Fig. 2**). Microscopic examination of kidney sections 14 days post infection (dpi) revealed more acute tubular necrosis in mice infected with *hlyA^++^* UTI89 than those infected with WT or Δ*hlyA* UPEC (**Fig. 1C,D**). Furthermore, analysis of kidney sections obtained 14 dpi revealed increased expression of the well-defined renal epithelial injury biomarker Kim-1^29^ in tubules of *hlyA^++^*-infected mice (**Fig. 1C,E**). Also, compared with WT UTI89- or *hlyA^++^*-infected mice, mice inoculated with Δ*hlyA:1* or Δ*hlyA:2* exhibited a significant decrease in circulating Kim-1 protein at 14 dpi (**Fig. 1F**). These results demonstrate that independent of bacterial burden, HlyA secretion by UPEC enhances renal epithelial damage during pyelonephritis.

**Figure 1.**
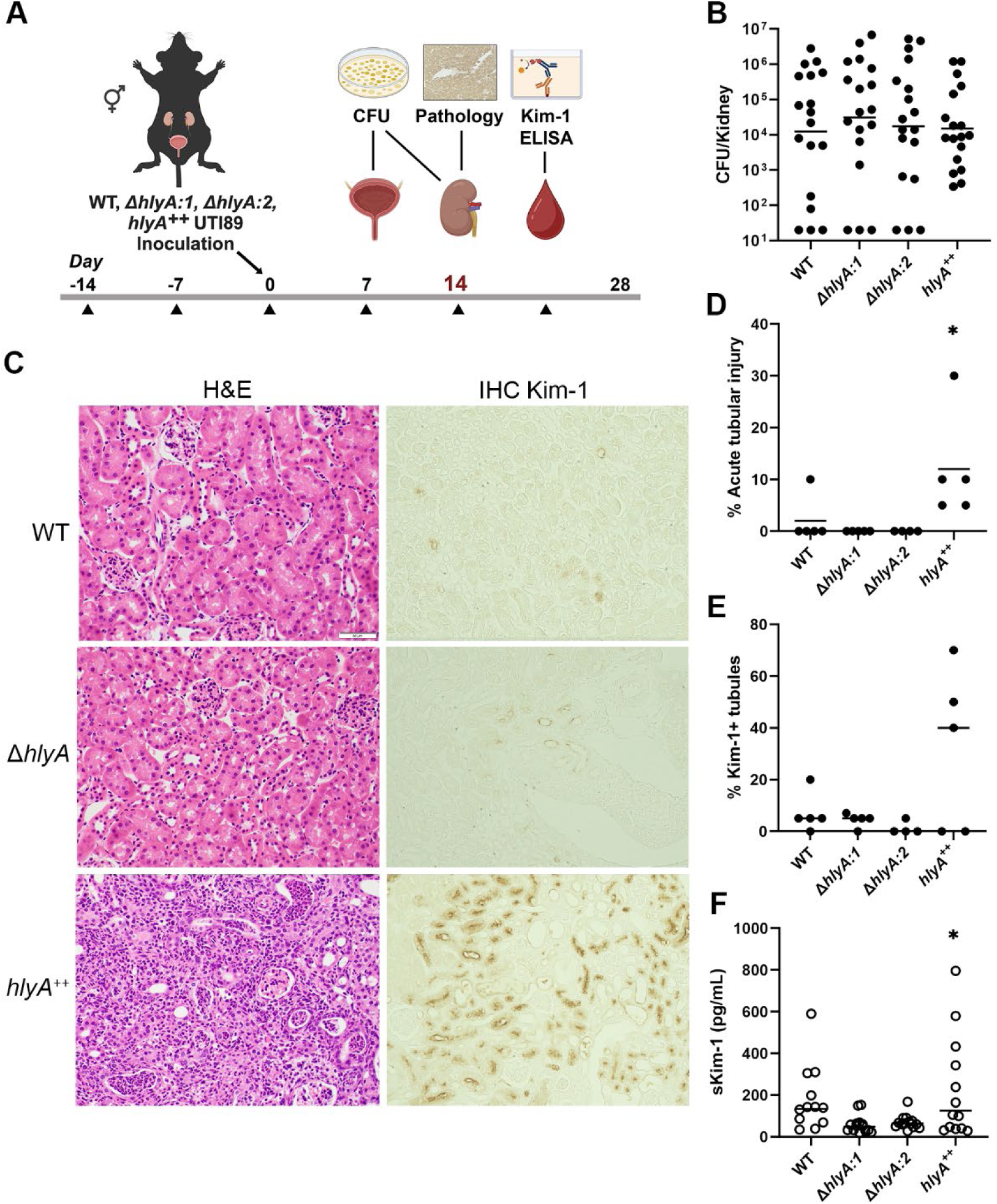
HlyA production exacerbates renal epithelial injury during pyelonephritis. **A,** Schematic of experimental procedures and timeline. Triangles indicate testosterone cypionate (or vehicle) dosing; all data in subsequent panels reflect 14 d post infection with the indicated UPEC strains. **B,** Bacterial loads (colony-forming units [CFU]) in the kidneys of infected mice (ANOVA not significant [ns]). **C,** Histology of mouse kidneys, as visualized by hematoxylin and eosin staining (H&E, left column) or immunohistochemistry for Kim-1 (IHC, right column). **D,** Proportion of kidney cortex affected by acute tubular necrosis (ATN). ANOVA p=0.016; *p=0.025 for *hlyA*^++^ vs Δ*hlyA*:1, p=0.035 for *hlyA*^++^ vs Δ*hlyA*:2 by Tukey post-hoc analysis. **E,** Proportion of cortical tubules that stain positivity for Kim-1. ANOVA p=0.043; p=0.064 for *hlyA*^++^ vs Δ*hlyA*:2 by Tukey post-hoc analysis. **F,** Serum Kim-1 measured by ELISA. ANOVA p=0.0082; *p=0.021 for *hlyA*^++^ vs Δ*hlyA*:1, p=0.034 for *hlyA*^++^ vs Δ*hlyA*:2 by Tukey post-hoc analysis.

### A genome-wide CRISPR-Cas9 screen identifies endocytosis as essential for HlyA toxicity to renal epithelial cells

Although *in vitro* studies have established that HlyA is cytotoxic to renal epithelial cells,^18,19,28^ an epithelial receptor for HlyA has not been identified. We grew UTI89 Δ*hlyA* transformed with the *hlyCABD* expression plasmid pSF4000^30^ in LB broth daily and filter-sterilized this medium prior to its prompt use in experiments (**Fig. 2A**). The HlyA content of this conditioned medium (HlyA-CM) across daily preparations was standardized by quantitative hemolysis of rabbit RBCs and expressed in arbitrary hemolytic units (HU). Importantly, given this modest daily variation in obtained HlyA concentrations, some figures reflect representative cell-culture experiments (each with several replicate wells) from among several experiments performed on different days. Among multiple cell lines tested, inner medullary collecting duct cells of the mouse kidney (IMCD-3) exhibited higher cytotoxicity upon HlyA-CM treatment than did HEK293T (human embryonic kidney fibroblast-like) cells, J774 (murine macrophage-like) cells, or C57BL/6 mouse bone marrow-derived macrophages (BMDMs) (**Fig. 2B**); of note, J774 and BMDMs express CD11a/CD18, the reported myeloid-cell receptor for HlyA.^21,22^ Cytotoxicity to any of these lines was low upon treatment with CM prepared identically from UTI89 Δ*hlyA* (**Supplementary Fig. 3A**).

**Figure 2.**
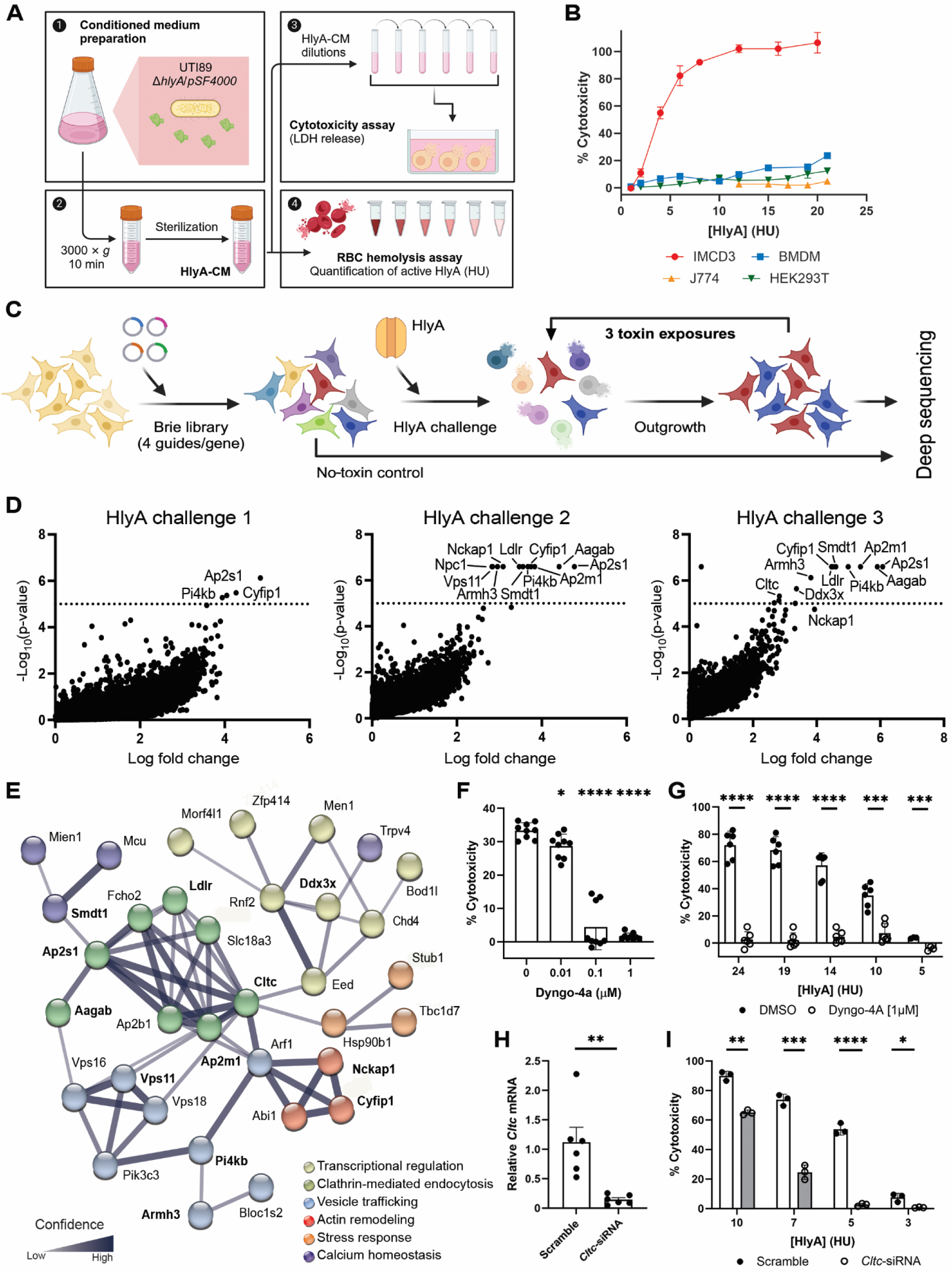
Host mediators of clathrin-mediated endocytosis confer renal epithelial susceptibility to HlyA. **A,** Schematic overview of HlyA-CM preparation. **B,** Percent cytotoxicity in IMCD3, BMDM, J774, and HEK293T cells exposed to HlyA-CM (1-22 HU) for 2 h. **C,** Experimental approach for the CRISPR-Cas9 screen. **D,** Volcano plots displaying log fold change in gene abundance versus -log_10_ p-value (dashed line = FDR threshold of 0.05) after sequential HlyA-CM challenges. **E,** String pathway analysis of the top 100 altered genes across the 3 toxin exposures; the 11 genes meeting FDR threshold are indicated in bold text. **F,** Percent cytotoxicity in IMCD-3 cells exposed to HlyA-CM (4 HU) for 2 h after pretreatment with Dyngo-4A or vehicle. ANOVA p<0.0001; *p=0.037, ****p<0.0001 vs control (0 Dyngo-4a) by Dunnett multiple comparison test. **G,** Percent cytotoxicity in primary C57BL/6 renal epithelial cells exposed to HlyA-CM for 2 h after pretreatment with Dyngo-4A or vehicle. ***p<0.001, ****p<0.0001 by unpaired t test. **H,** Relative *Cltc* expression in IMCD-3 cells 24 h after transfection with *Cltc*-siRNA or scramble control. **p<0.01 by unpaired t test. **I,** Percent cytotoxicity in IMCD-3 cells 24 h after transfection with *Cltc*<u>-</u>siRNA or scramble control, following HlyA-CM exposure for 2 h. *p<0.05, **p<0.01, ***p<0.001, ****p<0.0001 by unpaired t test. For all cytotoxicity data, individual experiments are shown and are representative of at least 3 biologically distinct experiments using CM produced on the day of each experiment.

Previously, we generated a CRISPR-Cas9-based gene disruption library in IMCD-3 cells,^23^ representing ∼4× nominal coverage of the entire mouse genome (**Supplementary Fig. 3B**). To identify host factors required for HlyA toxicity, a survival-based, positive selection screen was performed (**Fig. 2C**). The library of CRISPR-modified IMCD-3 cells was subjected to HlyA-CM (10 HU) for 2 h to produce ∼60% cell death; surviving cells were then expanded for 48 h. This exposure and recovery cycle was repeated 3 times, with surviving cells collected after each exposure and analyzed by deep sequencing to identify overrepresented sgRNA in surviving cells. The most significant gene “hits” over 3 toxin exposures were *Ap2s1*, *Aagab*, and *Ap2m1* (**Fig. 2D**), all of which encode components of the AP-2 adapter complex that links clathrin to the nascent membrane pit to initiate clathrin-mediated endocytosis. Other genes that reached statistical significance in the screen included additional ones participating in clathrin-mediated endocytosis, as well as the low-density lipoprotein receptor (*Ldlr*) (**Fig. 2D** and **Table 1**). String and GO PANTHER term analysis of the top 100 aggregate enriched genes over toxin exposures 2 and 3 showed strong relatedness in function and conserved roles in clathrin-mediated endocytosis and vesicle transport (**Fig. 2E** and **Supplementary Fig. 4**).

**Table 1.**
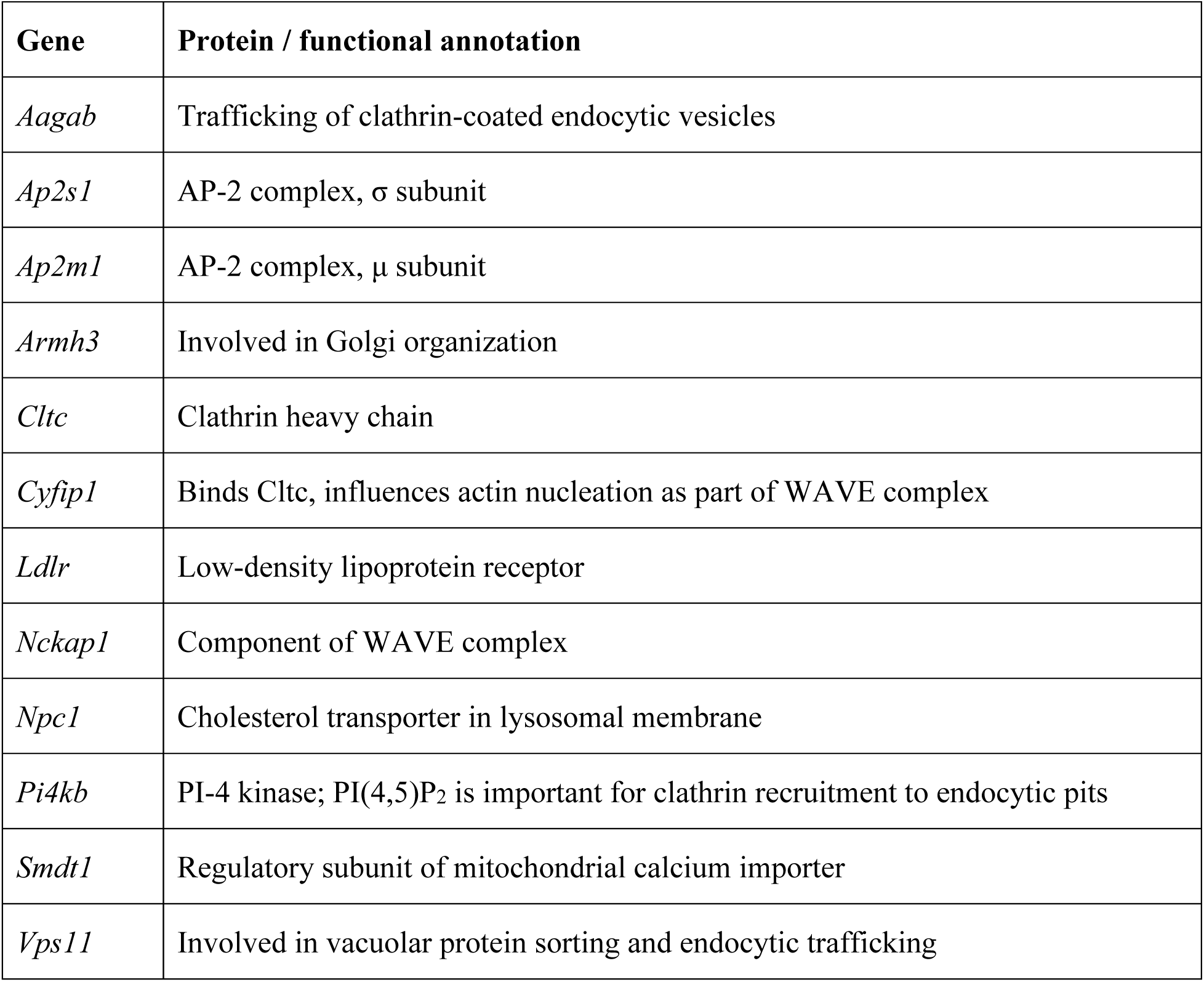
Statistically significant genes in the CRISPR mutant screen.

To validate a requirement for clathrin-mediated endocytosis in HlyA-mediated cytotoxicity to renal epithelial cells, we used pharmacological and genetic approaches. Dynamin-2, essential for scission of the invaginated membrane and separation of the endocytic vesicle, is inhibited by Dynasore^31^ and its derivatives, including Dyngo-4A. Pretreatment of IMCD-3 cells with Dyngo-4A before exposure to HlyA-CM reduced cytotoxicity in a dose-dependent manner, as compared to the vehicle (DMSO) control (**Fig. 2F**). We confirmed the known on-target effects of Dyngo-4A, as transferrin uptake was inhibited but total membrane cholesterol was unchanged (**Supplementary Fig. 5**). Pretreatment with Dyngo-4A also abolished HlyA-mediated cytotoxicity of primary renal epithelial cells from C57BL/6 mice (**Fig. 2G**). To exclude off-target effects of the dynamin inhibitor,^32^ we also conducted targeted downregulation of selected genes identified in the CRISPR screen with specific roles in clathrin-mediated endocytosis. siRNA-mediated downregulation of *Cltc* (clathrin heavy chain) (**Fig. 2H**) reduced the susceptibility of IMCD-3 cells to HlyA-mediated cell death, compared to the scramble siRNA control (**Fig. 2I**). Analogous results were seen with siRNA-mediated downregulation of *Ap2m1* (AP-2 complex, μ subunit; **Supplementary Fig. 6**).

### Internalized HlyA permeabilizes lysosomes and initiates necrotic cell death

Our genetic data suggested that HlyA internalization into epithelial cells is essential for its cytotoxic function, which contrasts with the hypothesis that HlyA lyses cell through pore formation in the plasma membrane.^14^ We speculated that endocytosed HlyA might porate lysosomes, enabling the release of lysosomal contents and ultimately necrosis. IMCD-3 cells treated with HlyA-CM and examined by immunofluorescence microscopy exhibited cellular rounding and nuclear condensation in a dose-dependent manner, whereas these changes were not seen in cells exposed to Δ*hlyA*-CM (**Supplementary Fig. 7A**). In contrast to cells treated with Δ*hlyA*-CM, those exposed to HlyA-CM after loading with LysoTracker Red dye exhibited a dose-dependent decrease in fluorescence, signifying lysosomal membrane permeabilization (**Fig. 3A**). HlyA-CM-exposed cells also exhibited a dose-dependent increase in pHrodo Green signal, reflecting cytoplasmic acidification, whereas cells exposed to Δ*hlyA*-CM did not show this change (**Fig. 3B** and **Supplementary Fig. 7B**). Cellular fractionation was performed to assess for the presence of lysosomal proteins in the cytoplasm of HlyA-CM-treated IMCD-3 cells. Immunoblotting demonstrated a dose-dependent increase in the lysosomal membrane protein LAMP-2 within the cytoplasm, compared with Δ*hlyA*-CM-treated cells (**Fig. 3C**). Within endolysosomal vesicles, multiple classes of proteases, including cathepsins, aid in the recycling or elimination of endocytosed proteins.^33^ Immunoblotting demonstrated a dose-dependent increase in the 46-kDa intermediate form of cathepsin D^34^ and the 32-kDa mature form of cathepsin D in the cytoplasmic fraction of cells exposed to HlyA-CM (**Fig. 3C**).

**Figure 3.**
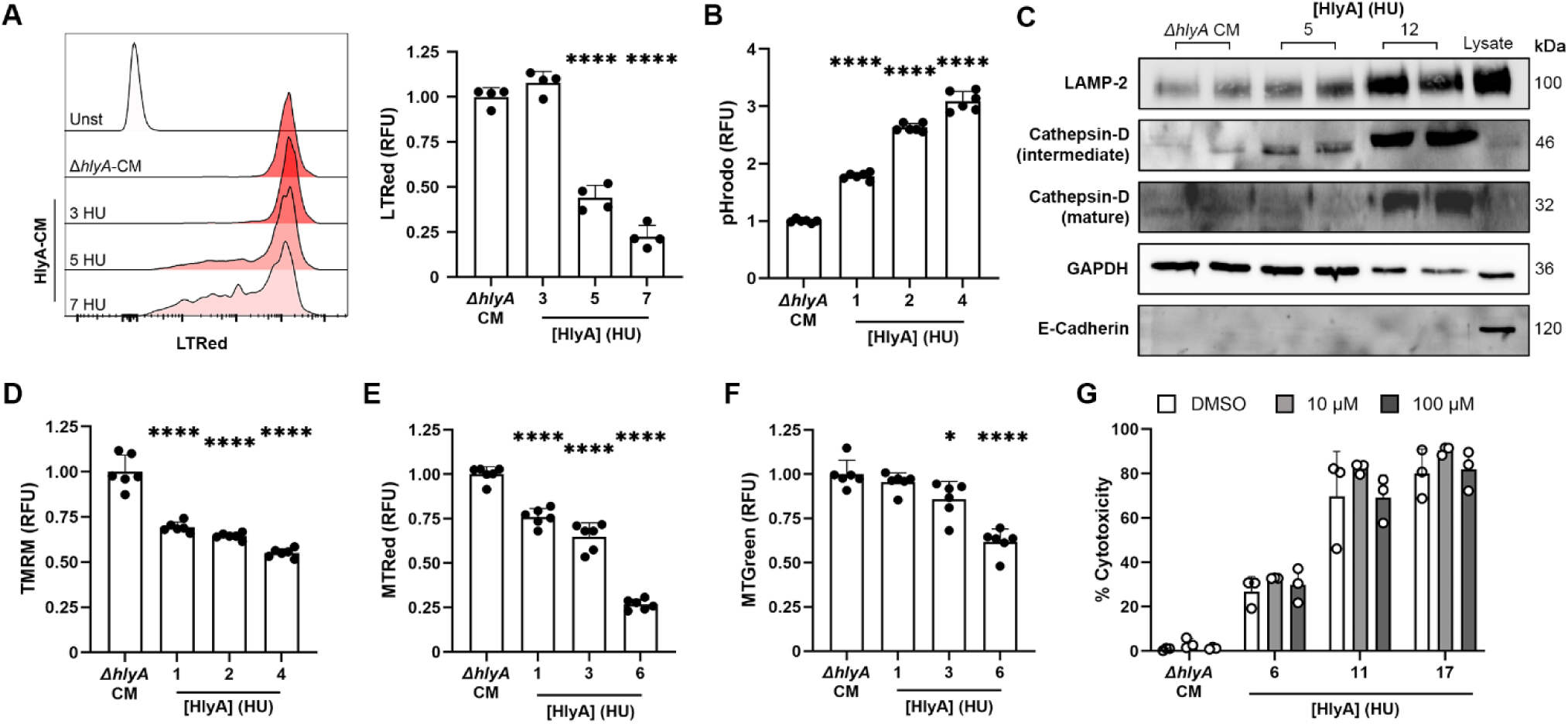
HlyA triggers lysosomal permeabilization and mitochondrial dysfunction preceding rapid caspase-independent cell death. **A,** Representative flow cytometry trace of LysoTracker Red signal (left) and quantification of lysosomal integrity (right) in IMCD-3 cells exposed to increasing concentrations of HlyA-CM or to Δ*hlyA*-CM. ANOVA p<0.0001; ****p<0.0001 vs Δ*hlyA*-CM by Dunnett multiple comparison test. **B,** Increasing pHrodo Green signal in IMCD-3 cells, reflecting decreased cytoplasmic pH, upon treatment with increasing concentrations of HlyA-CM or with Δ*hlyA*-CM. ANOVA p<0.0001; ****p<0.0001 vs Δ*hlyA*-CM by Dunnett multiple comparison test. **C,** Immunoblots for lysosomal proteins (LAMP-2, Cathepsin-D) in the cytoplasmic fraction of IMCD-3 cells exposed to HlyA-CM or Δ*hlyA*-CM. E-cadherin blot demonstrates no membrane contamination of the cytoplasmic fractions. **D-F,** Flow cytometry analyses of IMCD-3 cells 15 min post HlyA-CM or Δ*hlyA*-CM challenge. **D,** Mitochondrial membrane potential as measured by TMRM fluorescence. ANOVA p<0.0001; ****p<0.0001 vs Δ*hlyA*-CM by Dunnett multiple comparison test. **E,** MitoTracker Red fluorescence. ANOVA p<0.0001; ****p<0.0001 vs Δ*hlyA*-CM by Dunnett multiple comparison test. **F,** MitoTracker Green fluorescence. ANOVA p<0.0001; *p=0.014, ****p<0.0001 vs Δ*hlyA*-CM by Dunnett multiple comparison test. **G,** IMCD-3 cytotoxicity with HlyA-CM or Δ*hlyA*-CM exposure, after pre-treatment with the indicated concentrations of the pan-caspase inhibitor ZVAD-FMK; ANOVA ns. Fluorescence is expressed as geometric mean relative to fluorescence in the Δ*hlyA*-CM condition. For all cytotoxicity and flow cytometry data, individual experiments are shown and are representative of at least 3 biologically distinct experiments using CM produced on the day of each experiment.

Cathepsins and other lysosomal proteases in an acidified cytoplasm can cause degradation of mitochondrial membrane proteins, with ensuing mitochondrial dysfunction and cytosolic oxidation.^35^ In a recent study, HlyA diminished the mitochondrial membrane potential (ΔΨ_M_) and stimulated mitochondrial production of reactive oxygen species (ROS) in THP-1 macrophage-like cells.^36^ Exposure to HlyA-CM for 30 min in IMCD-3 cells loaded with the ΔΨ_M_ indicator dye tetramethylrhodamine (TMRM) resulted in decreased TMRM fluorescence, compared with Δ*hlyA*-CM treated cells (**Fig. 3D** and **Supplementary Fig. 7B**). Loss of mitochondrial membrane potential and mitochondrial mass was confirmed by diminished MitoTracker Red (**Fig. 3E**) and MitoTracker Green (**Fig. 3F**) signals in HlyA-CM-exposed cells (also see **Supplementary Fig. 7B**). Mitochondrial dysfunction can be part of several cell death pathways, and HlyA is reported to cause apoptotic or inflammatory cell death in various cell types.^17,19^ Here, pretreatment of IMCD-3 cells with the pan-caspase inhibitor Z-VAD-FMK had no impact on HlyA-mediated cell death (**Fig. 3G**), consistent with a model in which HlyA-induced lysosomal permeabilization leads to mitochondrial failure and cellular necrosis.

### LDLR expression facilitates HlyA toxicity to renal epithelium

IMCD-3 cells with disruption of *Ldlr* were enriched in our CRISPR-Cas9 screen, suggesting that expression of functional LDLR was required for HlyA-mediated cytotoxicity. Mammalian LDLR is a surface-exposed plasma membrane receptor for the low-density lipoprotein (LDL) particle, which has key roles in cholesterol homeostasis and the development of atherosclerosis.^37,38^ LDLR binds the ApoB protein component of the LDL particle, triggering clathrin-dependent endocytosis.^39,40^ LDLR also has been co-opted as a cellular receptor for certain viruses^41–46^ and *Clostridioides difficile* toxin A (TcdA) entry into cells.^46,47^ HEK293T cells express little to no LDLR^48^ and are insensitive to HlyA-mediated lysis (**Fig. 2B**). Co-transfection of HEK293T cells with human *LDLR* and the gene (*LRPAP1*) encoding the required chaperone RAP 24 h prior to HlyA-CM treatment conferred HlyA-CM sensitivity, compared with cells transfected with only *LRPAP1* (**Fig. 4A**). Expression of LDLR was confirmed by qRT-PCR and immunoblotting (**Fig. 4B,C**). Of note, membrane cholesterol content among these cell populations and conditions was unchanged (**Supplementary Fig. 8A**), excluding cholesterol alterations as a primary explanation for the findings.

**Figure 4.**
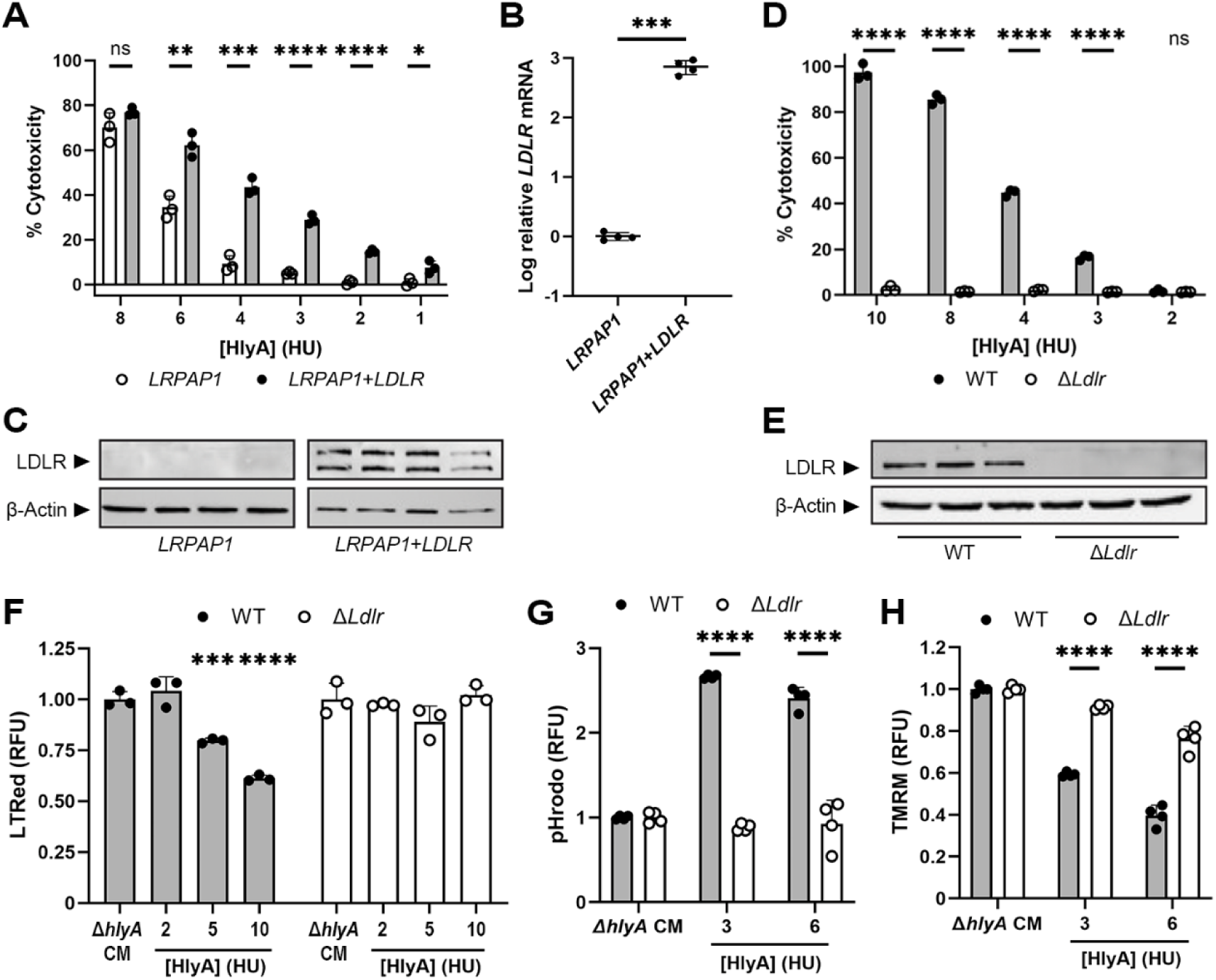
Low-density lipoprotein receptor (LDLR) expression enables HlyA cytotoxicity. **A,** Percent cytotoxicity in HEK293T cells following transfection with *RAP1* or *RAP1*+*LDLR* 24 h prior to HlyA-CM exposure. **B-C,** qPCR (**B**) and immunoblotting (**C**) for LDLR expression in HEK293T cells 24 h post-transfection with *LRPAP1* or *LRPAP1*+*LDLR*. **D,** Percent cytotoxicity in WT or Δ*Ldlr* IMCD-3 cells exposed to HlyA-CM for 2 h. **E,** Immunoblots demonstrating LDLR production in WT versus Δ*Ldlr* IMCD-3 cells. **F-H,** Flow cytometry-based analyses of WT or Δ*Ldlr* IMCD-3 cells challenged with HlyA-CM or Δ*hlyA*-CM as indicated. **F,** Lysosomal integrity as measured by LysoTracker Red fluorescence. ANOVA p<0.0001; ***p=0.008 and ****p<0.0001 vs Δ*hlyA*-CM by Dunnett multiple comparison test. **G,** cytoplasmic acidification as measured by pHrodo Green fluorescence. **H,** Mitochondrial membrane potential as measured by TMRM fluorescence. All fluorescence data are expressed as geometric mean relative to that in the Δ*hlyA*-CM condition. For all cytotoxicity and flow cytometry data, individual experiments are shown and are representative of at least 3 biologically distinct experiments using CM produced on the day of each experiment. RFU, relative fluorescence units. Comparisons in **A, B, D, G,** and **H** were made using unpaired t-tests, with *p<0.05, **p<0.01, ***p<0.001, ****p<0.0001.

In an orthogonal approach, targeted CRISPR-Cas9 editing of *Ldlr* was performed in IMCD-3 cells. Δ*Ldlr* IMCD-3 cells resisted HlyA-mediated killing over all HlyA-CM concentrations tested, whereas wild-type IMCD-3 cells were susceptible to HlyA-dependent cytolysis (**Fig. 4D**). We confirmed that Ldlr protein expression was absent in CRISPR-modified cells (**Fig. 4E**), with no impact on membrane cholesterol levels (**Supplementary Fig. 8B**). We confirmed the HlyA resistance phenotype in a second, independently generated IMCD-3 Δ*Ldlr* clone to exclude off-target gene editing effects (**Supplementary Fig. 9A**). IMCD-3 Δ*Ldlr* cells also did not exhibit the phenotypes of lysosomal damage, cytoplasmic acidification, or loss of mitochondrial membrane potential (**Fig. 4F-H** and **Supplementary Fig. 9B**) that were seen in WT cells upon exposure to HlyA-CM. Taken together, these data show that LDLR expression in human or mouse epithelial cells promotes HlyA-mediated cytotoxicity.

### Inhibition of LDLR-HlyA interaction protects renal epithelium from injury

To determine if HlyA binding to IM CD-3 cells depends on LDLR expression, we first studied whole-cell binding of HlyA using WT IMCD-3 or Δ*Ldlr* cells that were pretreated with Dyngo-4A to block CME (and therefore cytotoxicity) before HlyA-CM exposure. Supporting a function for LDLR as a receptor for HlyA, we found more HlyA bound to WT than Δ*Ldlr* IMCD-3 cells by immunoblotting lysates with an antibody recognizing HlyA (**Fig. 5A**). Furthermore, when physiologically relevant doses of purified human LDL (the canonical LDLR ligand) or a human APOB peptide (containing site B, the LDLR-binding moiety within LDL particles) were supplemented into serum-free HlyA-CM, dose-dependent reduction in cytotoxicity of WT IMCD-3 cells was observed, whereas supplementation with bovine serum albumin (as control) was not protective (**Fig. 5B,C**). Finally, emerging knowledge of LDLR-mediated viral entry has guided the development of protein-based molecules capable of protecting from infection with certain viruses.^49^ Accordingly, we tested whether addition of a recombinant fusion protein containing the human LDLR ectodomain (LDLR-Fc) could function as a decoy receptor to protect renal epithelial cells from HlyA action. Indeed, LDLR-Fc dose-dependently inhibited HlyA-induced toxicity toward IMCD-3 cells (**Fig. 5D**) and primary renal epithelial cells (**Fig. 5E**). Together, these results indicate that direct interaction between HlyA and LDLR precedes toxin internalization and cellular toxicity.

**Figure 5.**
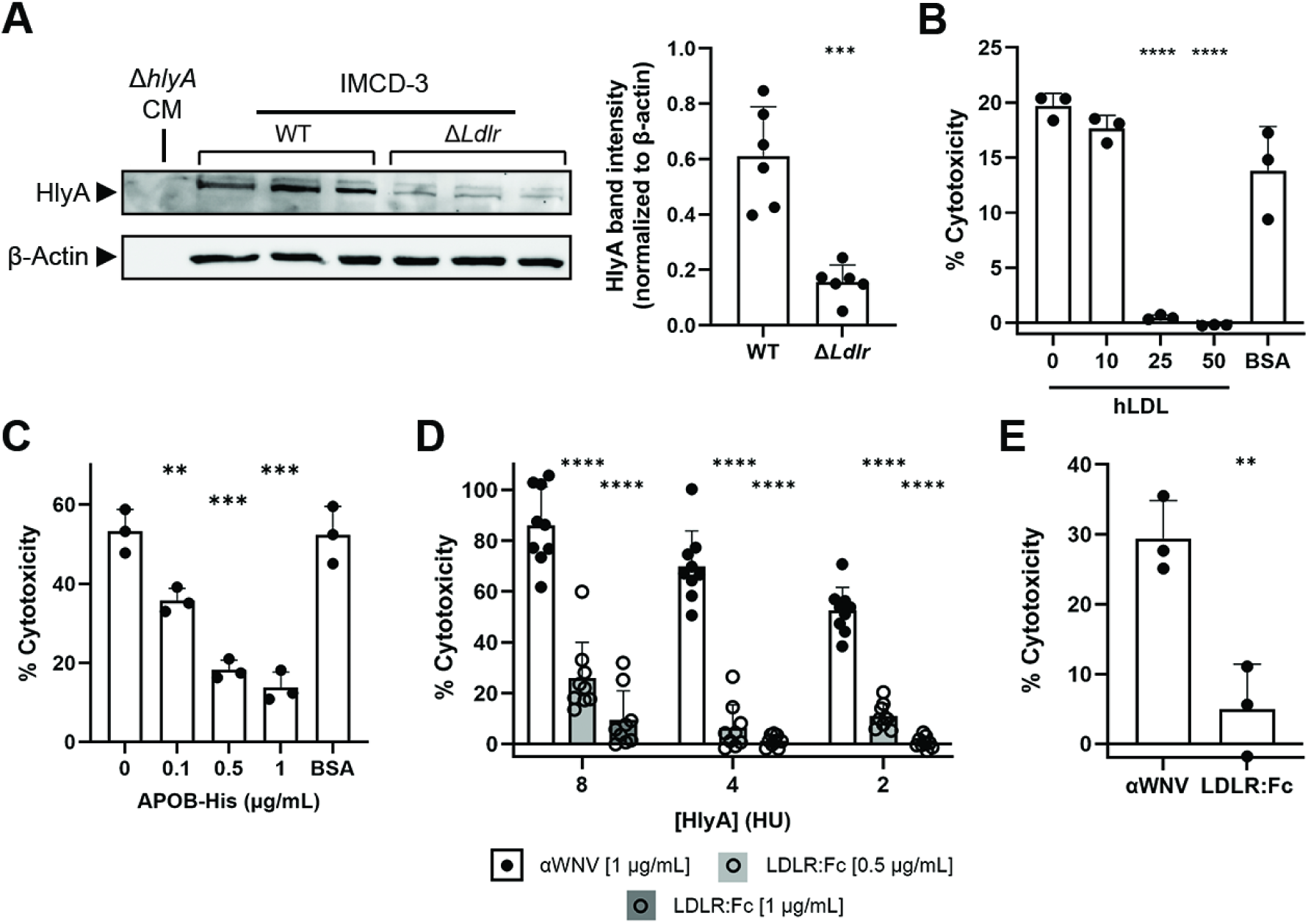
Targeting LDLR-HlyA interactions abrogates cytotoxicity to renal epithelial cells. **A,** Immunoblot (left) for HlyA on cell lysates after treatment of WT or Δ*Ldlr* IMCD-3 cells with HlyA-CM; quantification shown at right (***p<0.001; by unpaired t test). **B,** Percent cytotoxicity in WT or Δ*Ldlr* IMCD-3 cells exposed for 2 h to HlyA-CM (4 HU) with addition of human LDL (at the indicated concentrations) or BSA (control). ANOVA p<0.0001; ****p<0.0001 vs 0 LDL by Dunnett multiple comparison test. **C,** Percent cytotoxicity in WT or Δ*Ldlr* IMCD-3 cells exposed for 2 h to HlyA-CM (7.5 HU) after addition of APOB-His (at the indicated concentrations) or BSA (control). ANOVA p<0.0001; ****p<0.0001 vs 0 LDL by Dunnett multiple comparison test. **D,** Inhibition of HlyA-mediated toxicity to IMCD-3 cells upon treatment with the indicated concentrations of LDLR-Fc fusion protein, compared with an unrelated control antibody (anti-West Nile virus hE16 [αWNV]). At each HlyA-CM concentration, ANOVA p<0.0001, ****p<0.0001 vs αWNV control by Dunnett multiple comparison test. **E,** Similar inhibition of HlyA-mediated toxicity to primary murine renal epithelial cells upon treatment with LDLR-Fc (1 μg/mL); **p<0.01 by unpaired t test. For all cytotoxicity data, individual experiments are shown and are representative of at least 3 biologically distinct experiments using CM produced on the day of each experiment.

## DISCUSSION

The UPEC α-hemolysin (HlyA) can exert cytotoxicity in a range of mammalian cell types. In this study, we demonstrate that HlyA action on renal epithelium relies on clathrin-mediated endocytosis and LDLR expression, and that rapid cell death ensues from lysosomal permeabilization, cytoplasmic acidification, and resulting mitochondrial injury. This work reveals a distinct paradigm for the biological activity of HlyA, a prototypic member of the RTX family that has been studied for decades but whose mechanisms of action have been elusive.

Compared with other families of bacterial pore-forming toxins (PFTs), the RTX toxins – comprising over 1,000 members across a wide range of pathogenic species – have proven more enigmatic despite extensive study: structural determinations have been difficult, diversity among their functional domains is broad, and even how the canonical acylation step enables toxin function is not understood. HlyA was initially isolated based on its capacity to lyse RBCs. The traditional view that HlyA kills cells via poration of the plasma membrane is based on biochemical and membrane-conductance observations with RBCs and artificial membranes (reviewed in ^50^). Similarly, studies of myeloid cell susceptibility to HlyA-mediated killing have been consistent with this mode of action. Nonetheless, typical purification and structure elucidation has been precluded by the instability and thermolability of the toxin, which was recognized some 60 years ago.^13^ As a result, prior work with HlyA has relied in part on harsh purification and reconstitution protocols that might affect toxin activity. Here, using conditioned medium from an overexpressing UPEC strain prepared fresh daily, we observed that renal epithelial cells were more susceptible than myeloid cells to HlyA action, as observed in at least one other report.^51^ The apparent preference of HlyA for myeloid cells (particularly over RBCs) has been attributed to their expression of CD11a/CD18 (LFA-1). Most recently, screening of a mutant library in myeloid U937 cells exposed to HlyA identified CD18, the integrin β_2_ subunit, as the most prominent hit. CD18 was shown to bind HlyA and a related RTX toxin, *Aggregatibacter actinomycetemcomitans* RtxA, and siRNA-mediated downregulation of CD18 protected U937 cells from HlyA action.^22^ As epithelial cells do not express CD18, we sought alternative cellular mediators of HlyA toxicity using a genetic screen. We identified a requirement for LDLR expression and clathrin-mediated endocytosis for HlyA activity in epithelial cells. The present findings do not refute earlier proposed mechanisms of HlyA action but rather expand our understanding of the activities of this and potentially other RTX family members.

Clathrin-mediated endocytosis is a common form of receptor-dependent ligand internalization in mammalian cells. Clathrin-mediated endocytosis and LDLR family members have been implicated in the pathogenic activity of other microbial toxins, though from families other than RTX, including the large *Clostridium* toxin (LCT) family. For example, activity of *Clostridioides difficile* exotoxin TcdA is reduced in epithelial cells lacking LDLR, though direct TcdA-LDLR interaction was not observed, suggesting LDLR might have a facilitating role after initial TcdA binding to surface glycosaminoglycans.^47^ Similarly, several LDLR family members (including LDLR, LRP1, and megalin) act via their ligand-binding domains to promote internalization of *Clostridium novyi* alpha-toxin (Tcnα).^52,53^ LRP1 also serves as a receptor for the uniquely structured vacuolating cytotoxin (VacA) of *Helicobacter pylori*,^54^ the *Pseudomonas aeruginosa* exotoxin A (an ADP-ribosylating toxin),^55^ and the deaminase toxin of *Pasteurella multocida*.^56^ Anthrax toxin, an example of the A-B toxin class, gains entry to target cells via the LDLR family member LRP6.^57^ As noted earlier, LDLR family members are also co-opted for cellular entry by an array of viruses, including encephalitic alphaviruses,^41,42,45,49,58^ vesicular stomatitis virus,^43^ and hepatitis B virus.^44^ Our screen identified LDLR itself as a primary HlyA receptor on renal epithelial cells, but we anticipate that other LDLR family members might specifically or redundantly facilitate HlyA toxicity, given their sequence and structural relatedness and based on their relative expression by various cell types. Of note, among the multiple means of LDLR inhibition we utilized, ApoB and LDL, as natural ligands for LDLR, might additionally reduce HlyA toxicity by causing internalization of the receptor.

Cholesterol in target membranes is essential to the stabilization and functionality of many PFTs, including HlyA^59^ and a separate toxin family in Gram-positive bacteria, the cholesterol-dependent cytolysins (reviewed recently in ^60^). We verified that the results observed during our perturbations of clathrin-mediated endocytosis and LDLR expression and our competitive inhibition studies were not due to altered plasma membrane cholesterol content, which might have imparted a nonspecific effect on HlyA activity. Another significant hit in our CRISPR screen was *Npc1*, which encodes a well-characterized lysosomal cholesterol transporter. Cholesterol may be important for lysosomal permeabilization, one of the first events following cellular internalization of HlyA. We detected lysosomal proteases in the cytoplasm of HlyA-intoxicated cells, and the accompanying cytoplasmic acidification might promote the activity of these proteases in this compartment. This model aligns with an earlier study reporting that in bladder epithelial cells treated with sublytic concentrations of HlyA, cellular protease activity in the cytoplasmic compartment led to degradation of cytoskeletal proteins such as paxillin.^17^ We suspect but have yet to demonstrate HlyA pore formation in the lysosomal membrane. At least two recent studies have demonstrated that lysosomal protease activity in the cytoplasm causes degradation of mitochondrial proteins and impairment of mitochondrial membrane potential and function,^35,61^ matching the sequence of events we observed during the rapid HlyA-induced and caspase-independent death of renal epithelial cells.

Our findings introduce a distinct mechanism of action for HlyA, and potentially for other RTX toxins. We postulate that the specific effects of HlyA on a given target cell likely depend on receptor expression, other plasma membrane characteristics, and toxin dose. Ongoing work includes further specification of HlyA trafficking after internalization, interrogation of other LDLR family members as potential mediators of HlyA toxicity in other cells, and efforts to elucidate structural features of the HlyA interaction with LDLR. The present work also illuminates possible anti-virulence therapeutic avenues to prevent tissue injury during UPEC pyelonephritis and other infections caused by HlyA-secreting *E. coli*.

## METHODS

### Bacterial culture

Uropathogenic *Escherichia coli* strains utilized in this study were derived from the clinical isolate UTI89.^62^ For murine UTI and for preparation of HlyA conditioned medium (CM; see below), bacteria were grown statically at 37°C for 18 h in Luria-Bertani (LB) broth.

### Genetic manipulation of UTI89 *hlyA* expression

Targeted genetic deletion and chromosomal mediated overexpression of *hlyA* were achieved via the lambda red recombinase system.^63^ Oligonucleotides used are listed in **Supplementary Table 1**. For deletion of *hlyA,* a chloramphenicol resistance cassette (*Cm^R^*) was amplified by PCR using primers comprising FLP recombinase sequences and regions complementary to those directly up-and downstream of the *hlyA* gene. This *Cm^R^* linear fragment was transformed into UTI89 carrying a plasmid (pKM208) expressing the lambda red recombinase. Cm^R^ clones were next transformed with the FLP expression plasmid pCP20 to effect excision of the *Cm^R^* gene. To account for potential off-target effects, two independently derived isolates (Δ*hlyA:1,* Δ*hlyA:2*) were generated and tested. To induce stable HlyA overexpression in UTI89, the *Cm^R^* cassette was cloned directly upstream of the *hlyCABD* regulatory leader sequence.^9^ All clones were verified by sequencing, *hlyA* expression was confirmed by qPCR, and hemolytic activity was verified by plating on 5% sheep blood agar.

### Preparation of conditioned medium

CM were prepared daily and immediately prior to use in assays. As a result, modest daily variation in HlyA concentration (as measured by quantitative hemolysis; see below) was observed. Preparation of HlyA-CM utilized UTI89 Δ*hlyA:1* transformed with the HlyA expression plasmid pSF4000.^30^ Briefly, 3 × 10^7^ CFU/mL of Δ*hlyA:1*/pSF4000 or Δ*hlyA:1* from static overnight LB broth culture was inoculated into complete or serum-free DMEM:F12 medium (as indicated) and incubated statically at 37°C for 1 h. Bacteria were removed via centrifugation and passage through a 0.22-μm sterilizing filter. Multiple dilutions of HlyA-CM (in DMEM:F12) were used in the *in vitro* assays described below.

### HlyA activity quantification

Concurrent with use of daily HlyA-CM preparations in cytotoxicity studies (see below), toxin activity was quantified using a hemolysis assay as previously described^64,65^) with the following alterations. Pre-washed rabbit RBCs were added to dilutions of HlyA-CM to a final concentration of 5% (v/v) in 500 μL, and the mixtures were incubated at 37°C with gentle agitation for 1 h. Triton X-100 (0.1%) served as positive control. After centrifugation, the optical density of supernatants (540 nm) was quantified on a Tecan M-Plex microplate reader. We defined one hemolytic unit (HU) as the amount of toxin required to induce 50% lysis of RBCs, as has been classically defined.^66^ Linear regression was performed to quantify the activity (HU) present within each HlyA-CM preparation and dilution.

### Murine UTI

Female C57BL/6 mice (Envigo) were exposed to androgen as previously described,^67^ with 150 mg/kg testosterone cypionate (Depo-testosterone, Pfizer) via weekly intramuscular injection beginning at 5 weeks and continuing until sacrifice; control mice received similar injections of vehicle (cottonseed oil). Overnight UPEC cultures were centrifuged and resuspended to ∼5 × 10^8^ colony-forming units (CFU)/mL in PBS. At 8-9 weeks, mice were inoculated via transurethral catheterization with ∼2.5 × 10^7^ CFU of the indicated UPEC strain. At the listed time points, mice were euthanized and organs harvested for downstream analyses. Bacterial loads were determined by serial dilution and plating of whole organ homogenates on LB agar. H&E-stained sections were prepared by the Washington University Anatomic and Molecular Pathology core laboratory. Single sections were examined for signs of acute tubular injury and scored in 10% increments over the entire cortex by a blinded board-certified pathologist (J.P.G.). For Kim-1 immunohistochemistry (IHC), the left kidney was bisected longitudinally prior to fixation and paraffinization. 8-μm sections were mounted on glass slides. Antigen retrieval was performed by boiling in 10 mM sodium citrate. Sections were probed with goat anti-Kim1 (R&D Systems AF1817, 1:250) before incubation with secondary biotinylated mouse anti-goat IgG (1:250) (Invitrogen, 31730, 1:250) followed by streptavidin-HRP (Thermo-Scientific N504, 1:250). Sections were treated with DAB substrate (Pierce) and dehydrated prior to mounting with glass coverslips. Kim-1-stained slides were examined and scored as percentage of positive tubules (defined as tubular brush border staining) in the most involved region of tissue (percent positive tubules per total tubules in 400× region). For Kim-1 quantification in serum, blood was collected 14 days post infection (dpi) via cardiac puncture and serum separated by centrifugation. Kim-1 ELISA (R&D Systems) was performed according to the manufacturer’s instructions.

### Cytotoxicity assays

To quantify HlyA-induced toxicity *in vitro,* epithelial cells were seeded into 96- or 24-well plates 24 h prior to HlyA-CM challenge. On the day of each experiment, medium was aspirated, and cells washed twice with sterile PBS prior to addition of HlyA-CM dilutions or control media. Cells were incubated at 37°C for 2 h, and then supernatants were removed and analyzed for lactate dehydrogenase (LDH) content using a commercial kit (Promega) per the manufacturer’s instructions. Triton X-100 (0.1%) treatment served as a control representing maximal cell death (LDH release). For experiments with endocytosis inhibition, Dyngo-4A (Sigma 324413; in 0.1% DMSO) was incubated at the specified concentrations with cells in serum-free DMEM:F12, beginning 1 h prior to and maintained during HlyA-CM challenge; 0.1% DMSO was included as vehicle control. In other experiments, recombinant LDLR ectodomain fusion peptide (LDLR-Fc; ACRO Biosystems), an unrelated control antibody (anti-West Nile virus hE16), purified human low-density lipoprotein (LDL) cholesterol (ThermoFisher #L3486), human APOB-His (LS Bio #LS-G14619), or bovine serum albumin (BSA, as control; ThermoFisher #BP9703) was added as specified, concurrent with HlyA-CM challenge. For all *in vitro* assays reported, data shown are representative of at least 3 independent experiments except where noted in specific Supplementary figures.

### CRISPR-Cas9 genetic screen

We previously reported the generation of a library of CRISPR-Cas9-modified murine IMCD-3 cells^23^ transduced with the Brie sgRNA library, representing roughly ∼20,000 mouse genes (4 sgRNAs per gene). Guide coverage of a freshly expanded aliquot of these cells was confirmed by sequencing. 5 × 10^6^ cells were seeded into 175-cm^2^ flasks in complete DMEM:F12 containing puromycin and blasticidin; 24 h after seeding, monolayers were exposed to ∼10-12 HU of HlyA-CM at 37°C for 2 h, visually achieving ∼60-80% cell death in the first round. The flask was gently washed with Dulbecco’s PBS (DPBS), medium was replaced with fresh complete DMEM:F12, and surviving cells were allowed to recover for 48 h. After recovery, cells were split 1:2 into new flasks; one flask proceeded to the next round of toxin exposure, while the other was saved for genomic DNA extraction. Sequencing was performed by the Genome Technology Access Center (GTAC@MGI) at Washington University, using the MAGeCK package (version 0.5.9)^68^ to generate read count tables for each sample and comparing conditions using the RRA algorithm. Candidate genes considered significant were those exhibiting average log fold change > 2 and false discovery rate <1%.

### CRISPR-mediated *Ldlr* disruption

*Ldlr* KO lines were created by the Genome Engineering & Stem Cell Center (GESC@MGI) at Washington University. Briefly, synthetic gRNA targeting the sequence 5’- CAGGAATGCATCGGCTGACANGG -3’ was purchased from IDT, complexed with Cas9 recombinant protein, and transfected into IMCD-3 cells. Transfected cells were then single-cell sorted into 96-well plates, and clones were sequenced to identify the indels desired for gene disruption. Two independent Δ*Ldlr* clones were generated and studied.

### Transferrin uptake

Cells were seeded at a density of 1.5 × 10^5^ cells per well in 24-well plates. The following day, medium was removed, and wells were washed with PBS before adding warm serum-free medium. Plates were incubated at 37°C with 5% CO_2_ for 30 min. Medium was then replaced with warm serum-free medium containing Dyngo-4A or DMSO (negative control) and incubated for an additional 30 min at 37°C with 5% CO_2_. Subsequently, fresh cold serum-free medium containing 25 µg/mL of AlexaFluor 488-labeled transferrin (JacksonImmuno 015-540-050) and either Dyngo-4A or DMSO (control) were added, and plates were incubated on ice for 10 min in the dark. Plates were transferred to a 37°C incubator for the indicated time intervals, after which they were promptly returned to ice. Medium was removed, and wells were washed twice with ice-cold PBS, twice with 500 µL ice-cold acid wash buffer, and twice more with ice-cold PBS. Next, 200 µL of 0.25% trypsin was added, and cells were incubated at room temperature for 3 min; 1 mL of PBS was added, and cells were transferred to FACS tubes. Cells were centrifuged and washed twice with PBS, then filtered into new FACS tubes; just prior to analysis, 200 ng of propidium iodide (PI) was added. Cells were analyzed on a Cytek Aurora instrument, and data were analyzed with FlowJo software.

### Lysosomal and mitochondrial assays

IMCD-3 cells (1.5 × 10^5^ per well) were seeded in 24-well plates with complete DMEM-F12 for 24 h. Cells were washed with PBS, then incubated with one of the following: 50 nM LysoTracker Red (ThermoFisher), pHrodo Green (ThermoFisher), MitoTracker Red, or MitoTracker Green, in serum-free DMEM:F12 for 30 min at 37°C. Cells were washed twice with PBS to remove excess dye and then exposed to HlyA-CM for 30 min at 37°C. Cells were liberated with trypsin as described above and washed twice with PBS in FACS tubes; PI was added immediately prior to analysis.

### Membrane cholesterol content

IMCD-3 monolayers grown to 80-90% confluency in 6-well plates were serum-starved for 30 min prior to incubation with Dyngo-4A or DMSO control in serum-free DMEM:F12. HEK293T cells transfected with *LRPAP1* or *LRPAP1*+*LDLR* were grown to 80-90% confluency in 6-well plates. In both cases, cells were liberated with 0.25% trypsin and washed with PBS prior to analysis with a commercial assay kit (Promega #J3190) according to the manufacturer’s instructions.

### Protein extraction, cellular fractionation, and immunoblotting

To measure LDLR expression, cells in 24-well plates were washed 3 times with 0.5 mL of cold PBS, then lysed with 0.5 mL of 1X RIPA buffer + Halt Protease Inhibitor Cocktail (ThermoFisher). For HlyA whole-cell binding analysis, IMCD-3 cells grown to 80-90% confluency in T25 flasks were lysed with 0.2 mL 1X RIPA buffer + Halt Protease Inhibitor Cocktail and centrifuged at 16,000 × *g* for 5 min at 4°C to remove debris. For isolation of cytoplasmic fractions, IMCD-3 cells were grown to 80-90% confluency in T75 flasks. Cells were exposed to indicated amounts of HlyA-CM or Δ*hlyA*-CM for 30 min at 37°C. Monolayers were washed with cold PBS, and cytoplasmic fractions were extracted with a commercial kit (ThermoFisher #78840). Protein concentration in all cellular extracts were quantified by BCA assay (Pierce).

For immunoblotting of cell lysates, 40 μg of lysate was loaded per well and separated on polyacrylamide gels. After transfer to nitrocellulose membranes, lysates were probed overnight with rabbit anti-LDLR (Invitrogen #MA5-32075, 1:1000), Rabbit anti-β-actin (Cell Signaling Tech #4967, 1:20,000), or mouse anti-HlyA^69^ (H10, 1:2000). Blots were incubated with the appropriate secondary antibodies for 1 h: donkey anti-rabbit-HRP (Cytiva #NA934V, 1:5000) or sheep anti-mouse-HRP (Cytiva #NA931V, 1:5000). Blots were visualized using western ECL substrate (Bio-Rad #1705060) and imaged on a Bio-Rad ChemiDoc MP instrument. For HlyA whole-cell binding analysis, HlyA band intensities were quantified via ImageJ (Fiji 2) and normalized to β-actin.

### Transfection

Expression vectors for *LRPAP1* and *LDLR* were previously reported.^49^ Plasmid DNA was extracted from 250-mL LB broth cultures using Qiagen Plasmid Maxi Kit (Qiagen #12163). HEK293T cells grown to ∼75% confluency in a T75 flask were transfected with 30 μL of Lipofectamine LTX DNA Transfection Reagent, 10 μL of PLUS Reagent (Life Technologies #15338-100), and 5 μg of *LRPAP1* plasmid DNA with or without 10 μg of *LDLR* plasmid DNA. Approximately 18 h later, cells were plated (2 × 10^5^ cells per well in 24-well plates); 24 h after seeding, cells were exposed to HlyA-CM or harvested for LDLR immunoblotting.

### siRNA treatment

IMCD-3 cells were seeded in 24-well plates (∼10^5^ cells/well); 18 h later, cells were transfected with 50 nM of siRNA and 1 μL of Lipofectamine 2000 (ThermoFisher #11668027) per well following the manufacturer’s instructions. One day post-transfection, cells were used for cytotoxicity assays or RNA extraction. The fluorescent reporter TYE563 and housekeeping gene *Hprt* were used as positive controls to verify transfection, and scramble siRNA was used as control for *Cltc* and *Ap2m1* siRNA knockdown.

### RNA extraction and qPCR

Cells were grown to ∼80-90% confluency in 24-well plates, medium was removed and cells washed with PBS. RNA was isolated with RNA STAT-60 (Tel-Test Inc.) and cDNA synthesized (BioRad iScript) per respective manufacturer’s protocols. qPCR was performed using SYBR Green Supermix (BioRad). Each reaction contained 10 µL of SYBR Green master mix, 20 ng cDNA, and 0.5 µM forward and reverse primers (**Supplementary Table 1**) in a final volume of 20 µL.

### Statistical analysis

Pairwise comparisons of *in vivo* bacterial titers and renal pathology were made using the nonparametric Mann-Whitney U test. Other pairwise comparisons were made with unpaired t tests. Multiple comparisons were made using ANOVA with Tukey post-hoc or Dunnett multiple comparison tests as indicated. *p* values ≤ 0.05 were considered significant. All numerical data were analyzed using GraphPad Prism 10.4.

## ACKNOWLEDGMENTS

This work was supported by National Institutes of Health grants T32-007067 and F31-AI176711 (to H.W.K.); and R01-DK126697 and R01-AI158418 (to D.A.H.). We acknowledge Drs. Sean P.J. Whelan, Paul Rothlauf, and Stephen Sykes for helpful advice; Drs. Monica Sentmanat and Yong Miao of the Wash U Genome Engineering and Stem Cell Center (GESC@MGI) for CRISPR screen sequencing, data analysis, and creation of the IMCD-3 Δ*Ldlr* cell lines; Dr. Wandy Beatty for assistance with microscopy; and Dr. Rodney Welch for providing antisera recognizing HlyA. Figures 1A, 2A, 2C, and S1A were generated with Biorender.com.

## AUTHOR CONTRIBUTIONS

H.W.K., M.R.S., C.A.C., J.P.G., M.S.D., and D.A.H. designed experiments. H.W.K, M.R.S., R.J.J., C.A.C., H.M., and J.S. performed experiments and analyses; J.P.G. reviewed renal pathology. M.S.D. contributed LDLR-related reagents and constructs. H.W.K., M.R.S., and D.A.H. drafted the manuscript, and final editing was done by M.S.D. and D.A.H.

