## Supplementary Data for "LDL receptor-mediated endocytosis of *Escherichia coli* α-hemolysin mediates renal epithelial toxicity"

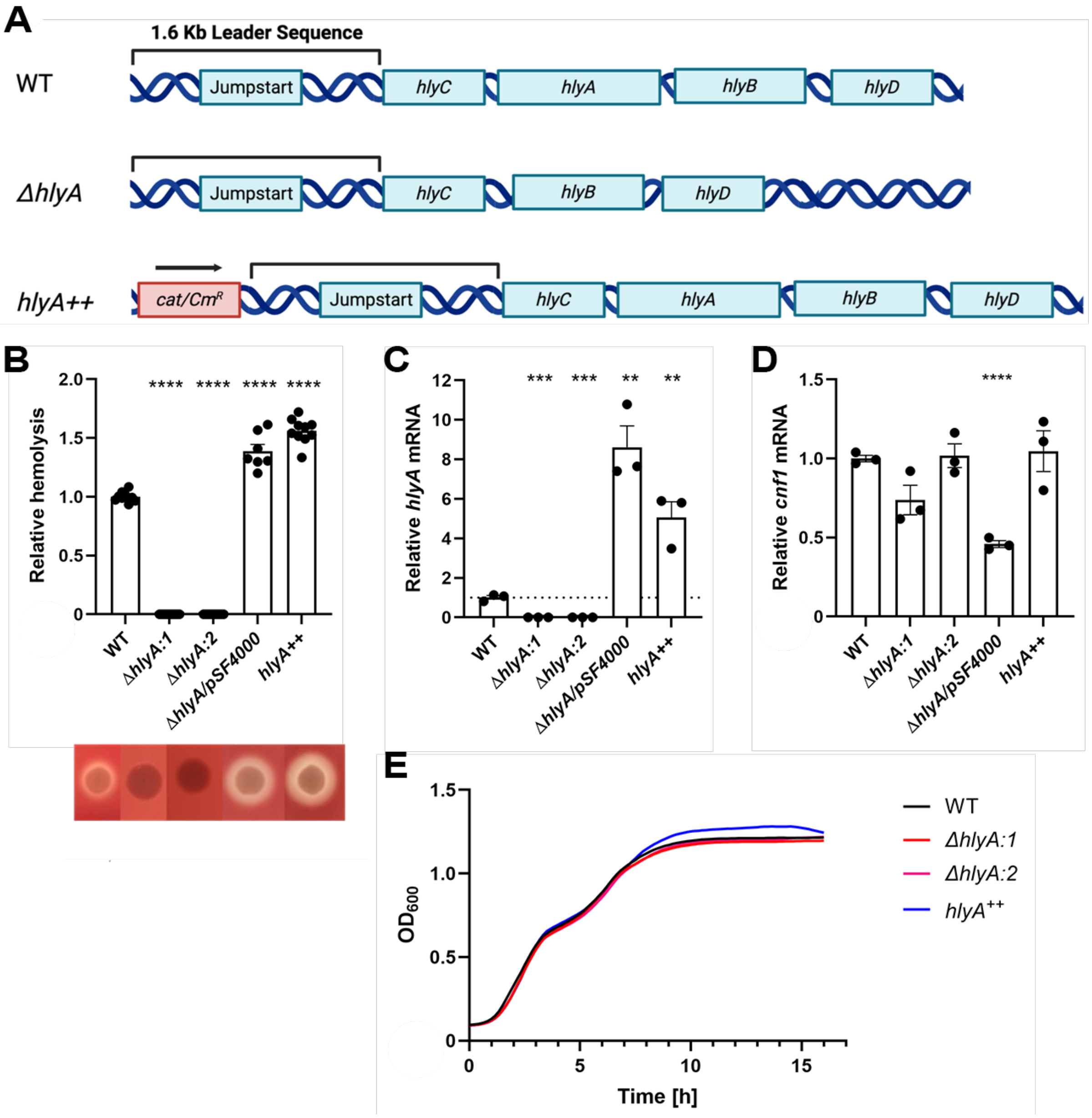


**Supplementary Figure 1. Construction and phenotypic characterization of UPEC mutants with differing HlyA secretion. A,** Schematic representation of design of UTI89 strains with differential HlyA secretion capacity, including two independent Δ*hlyA* clones. **B,** Quantification of hemolytic activity of engineered UTI89 HlyA secretion mutants on 5% sheep blood agar (representative images below the graph). **C,** qPCR of *hlyA* expression by the indicated strains. **D,** qPCR of *cnf1* expression (downstream of the *hlyCABD* operon) by the indicated strains. **E,** Growth curves of the strains in LB broth (point symbols omitted for clarity). **p<0.01, ***p<0.001, ****p<0.0001 by ANOVA with Dunnett multiple comparison test.

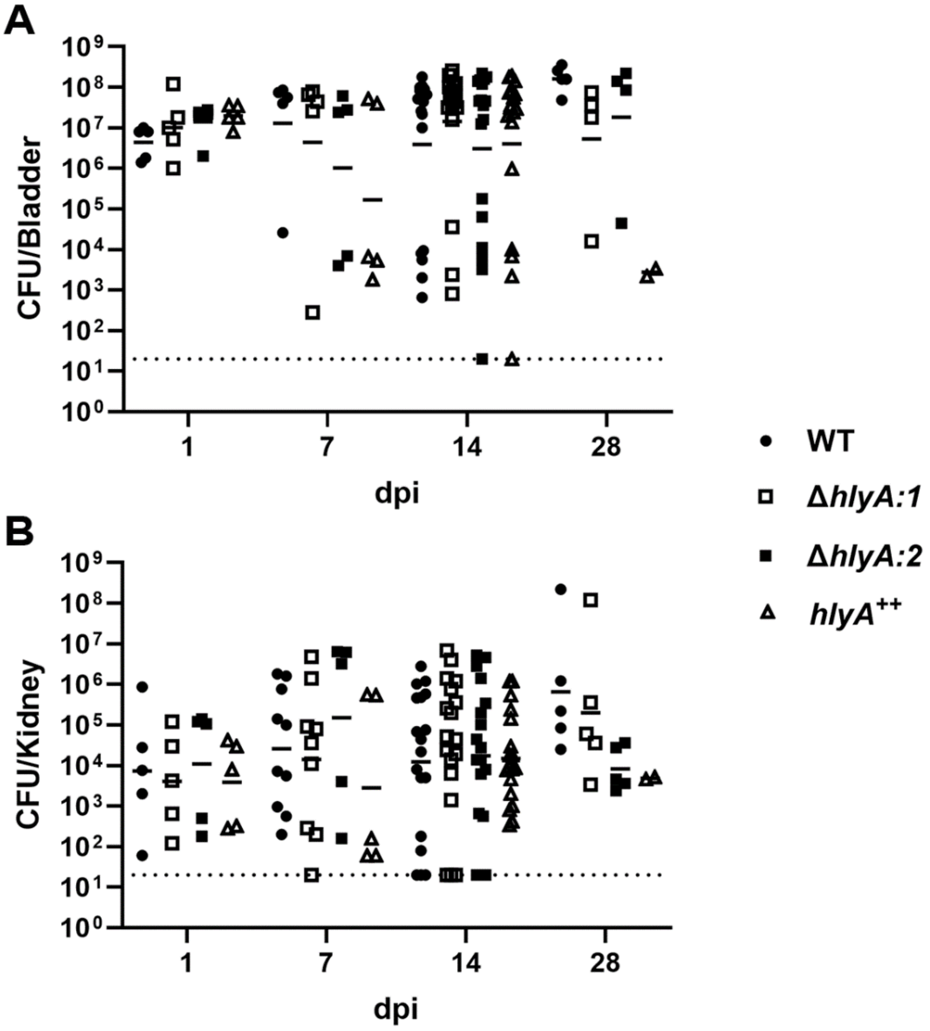


**Supplementary Figure 2.** Bacterial loads in the bladder **(A)** and kidney **(B)** at the indicated days post infection (dpi) following UPEC infection of androgen-exposed female C57BL/6 mice. Day 14 kidney bacterial loads are the same as shown in **Fig. 1B**. CFU, colony-forming units. n = 5-18 mice per condition at each time point; no significant differences between infecting strains by ANOVA.

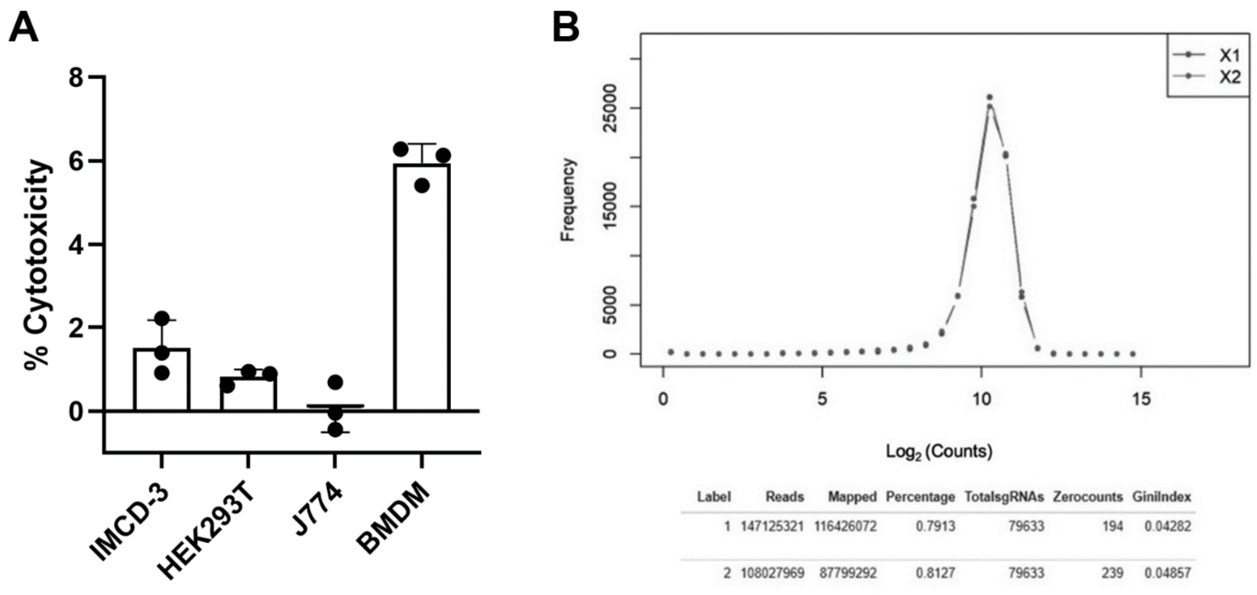


| Label | Reads | Mapped | Percentage | TotalsgRNAs | Zerocounts | GiniIndex |
|---|---|---|---|---|---|---|
| 1 | 147125321 | 116426072 | 0.7913 | 79633 | 194 | 0.04282 |
| 2 | 108027969 | 87799292 | 0.8127 | 79633 | 239 | 0.04857 |

**Supplementary Figure 3. A,** Minimal cytotoxicity of Δ*hlyA*-CM to various cell lines (note y-axis range). **B,** Verification of CRISPR mutant library coverage (79,633 gRNAs).

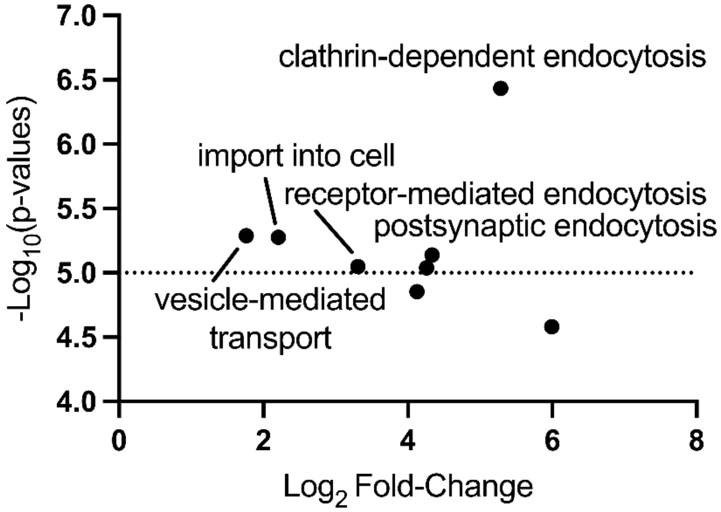


**Supplementary Figure 4.** GO PANTHER pathway terms representing the guides enriched in surviving CRISPR-modified IMCD-3 cells.

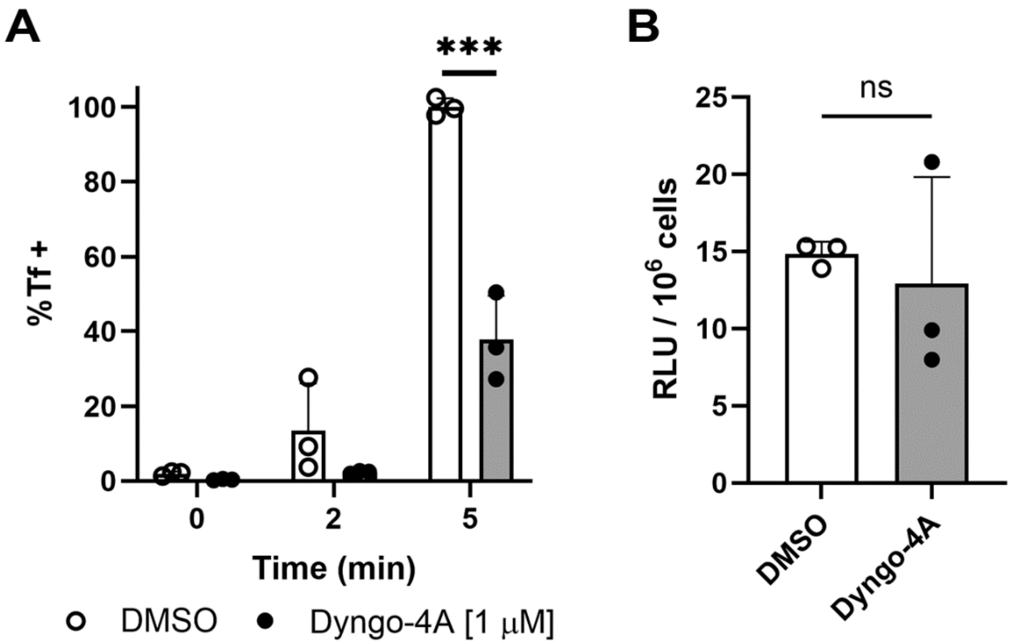


**Supplementary Figure 5.** Fluorescent transferrin (Tf) uptake **(A)** and membrane cholesterol **(B)** in IMCD-3 cells treated with the dynamin inhibitor Dyngo-4A or with DMSO control. Single experiment with technical replicates shown. RLU, relative luminescence units. ***p<0.001 by unpaired t test.

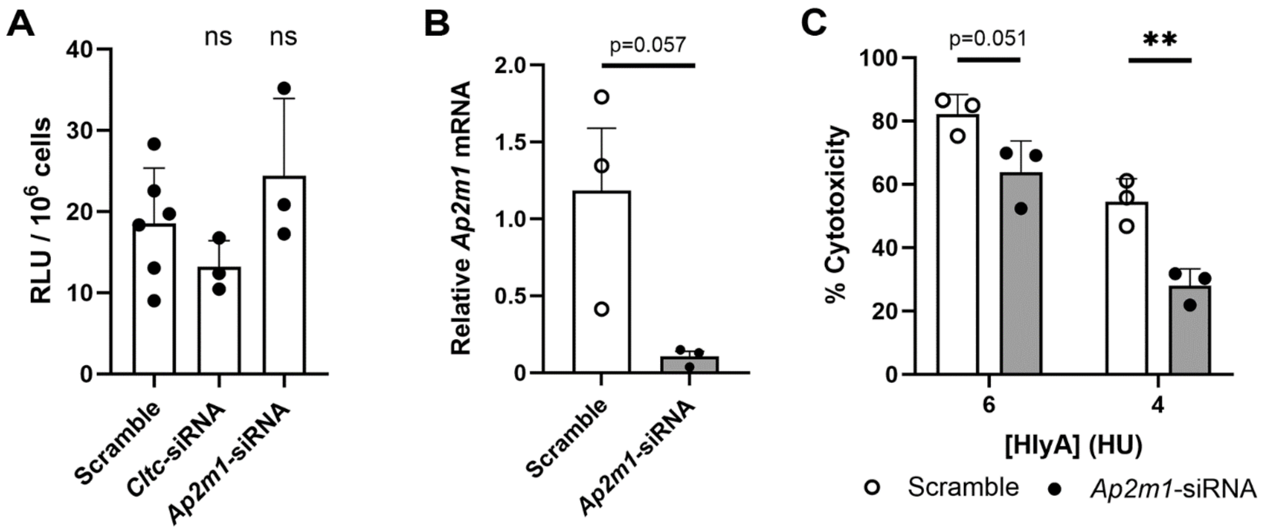


**Supplementary Figure 6. A,** Membrane cholesterol levels in IMCD-3 cells receiving siRNA targeting *Cltc* or *Ap2m1*, or scramble siRNA control (single experiment with technical replicates; ns by ANOVA vs Scramble). RLU, relative luminescence units. **B,** Knockdown of *Ap2m1* mRNA, measured by qPCR; p value as shown by unpaired t test. **C,** Reduced HlyA cytotoxicity to IMCD-3 cells with siRNA knockdown of *Ap2m1*. p values as shown and **p<0.01 by unpaired t test.

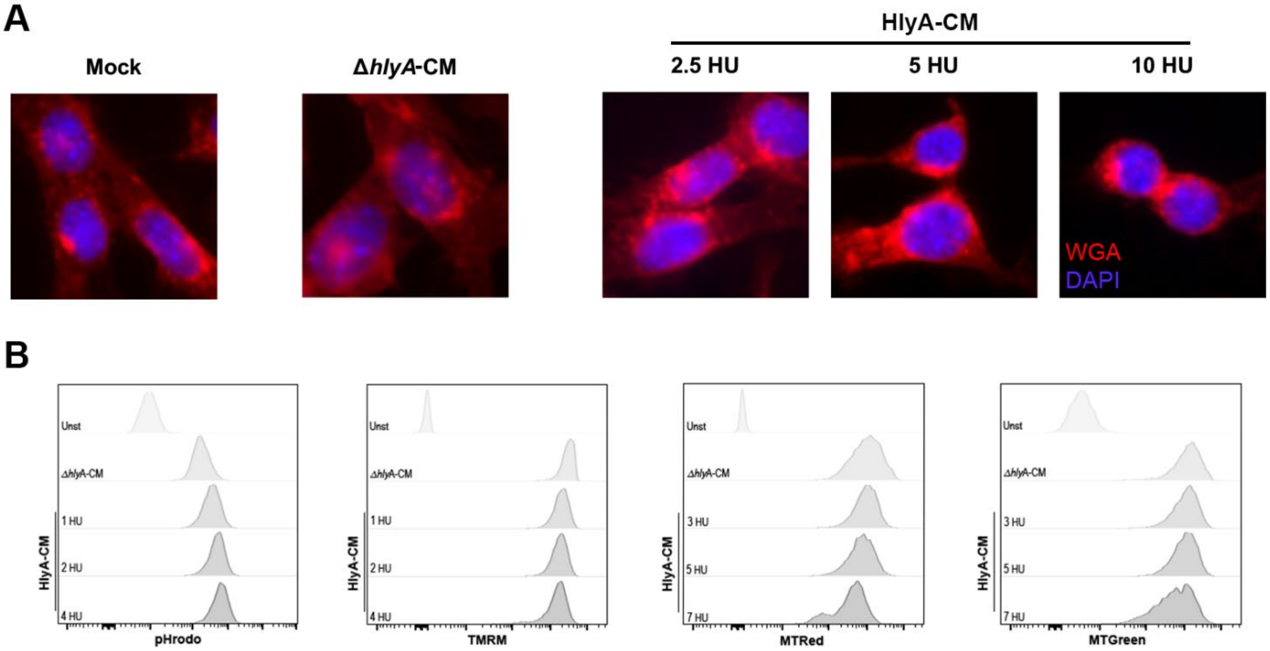


**Supplementary Figure 7. A,** Representative immunofluorescence microscopy images of cells exposed to Δ*hlyA-C*M or 2.5-10 HU of HlyA-CM, stained with wheat germ agglutinin (WGA; red) and DAPI (blue). **B,** representative histograms of IMCD-3 cells treated with HlyA-CM and analyzed by flow cytometry for pHrodo Green (see **Fig. 3B**), TMRM (**Fig. 3D**), MitoTracker Red (**Fig. 3E**), and MitoTracker Green (**Fig. 3F**).

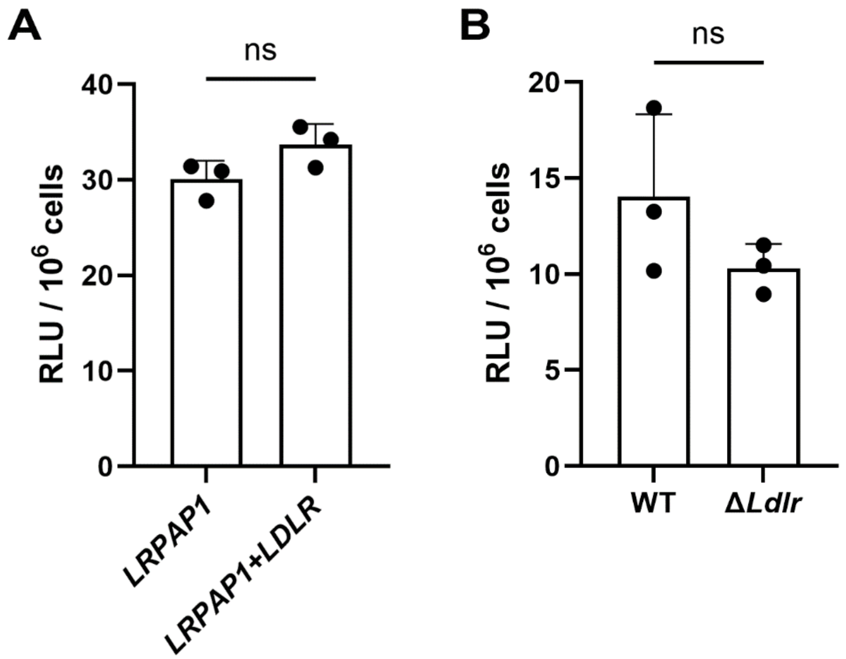


**Supplementary Figure 8. A,** Membrane cholesterol levels in HEK293T cells after transfection with *LRPAP1* or *LRPAP1* + *LDLR*. **B,** Membrane cholesterol levels in wild-type (WT) IMCD-3 cells and cells with CRISPR-mediated disruption of *Ldlr*. Single experiment with technical replicates shown. RLU, relative luminescence units.

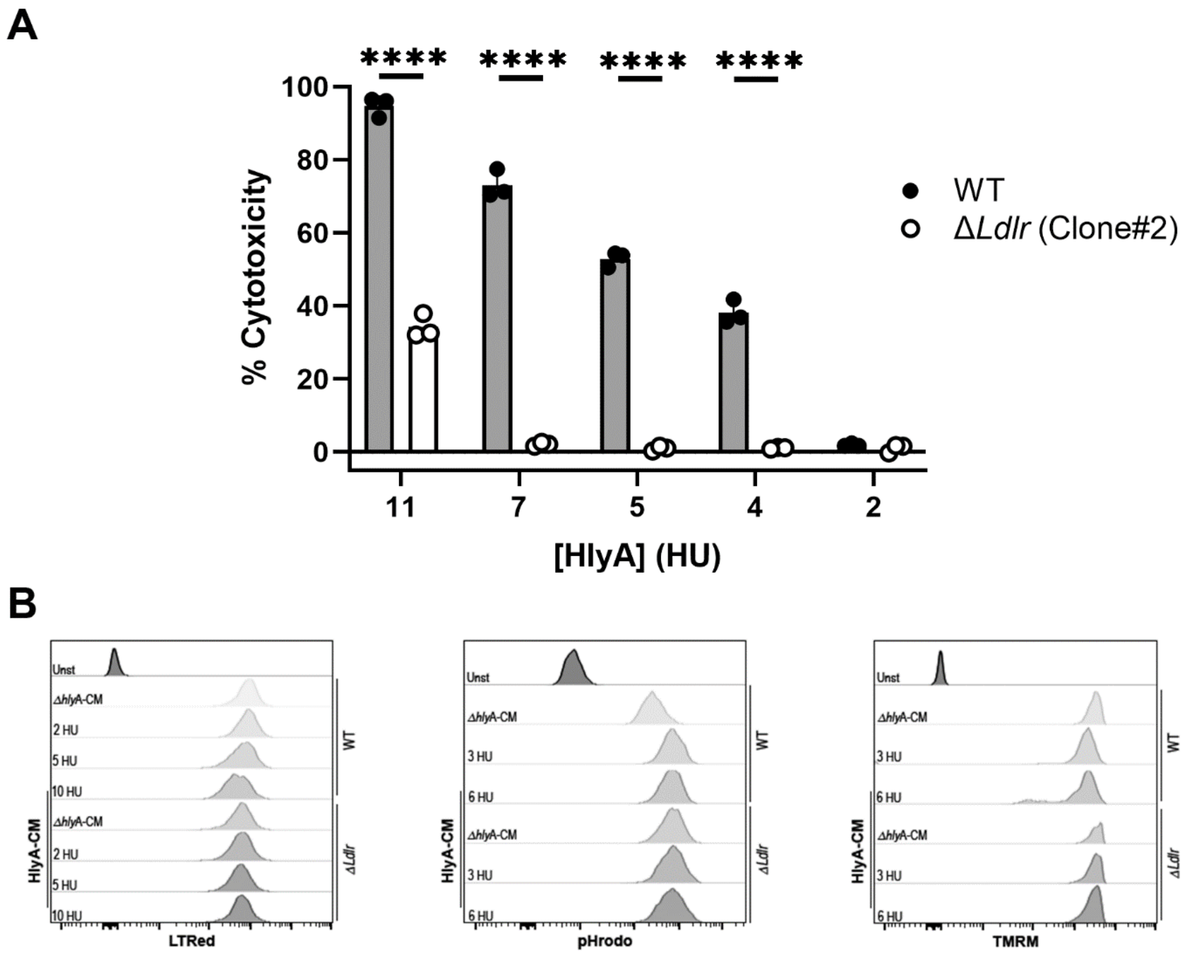


**Supplementary Figure 9. A,** CRISPR disruption of *Ldlr* in a second, independent clone of IMCD-3 cells confirms protection from HlyA-mediated cytotoxicity. ****p<0.0001 by unpaired t test. **B,** representative histograms of WT (upper traces) and Δ*Ldlr* (lower traces) IMCD-3 cells treated with HlyA-CM and analyzed by flow cytometry for LysoTracker Red (see **Fig. 4F**), pHrodo Green (**Fig. 4G**), and TMRM (**Fig. 4H**).

**Supplementary Table 1. Oligonucleotides used in this study.**

| Name | Sequence (5'-3') | Application |
|---|---|---|
| Δ*hlyA-Cm*$^R$ 5' | AATCTTATGTGGCACAGCCCAGTAAGATTGCTATCATTTAAATTAATATATGAAGTTCCTATACTTTCTAGAGAATAGGAACTTCGTGTAGGCTGGAGCTGCTTC | Genetic knockout |
| Δ*hlyA-Cm*$^R$ 3' | AAAAACAAGACAGATTTCAATTTTTCATTAACAGGTTAAGAGGTAATTAAGAAGTTCCTATTCTCTAGAAAGTATAGGAACTTCAATGGGAATTAGCCATGGTCC | Genetic knockout |
| *hlyA* Seq 5' | CGTATAACCCATAATCAATTTTATGACAAGAATCC | Sequence verification |
| *hlyA* Seq 3' | CACGGAGGTAAAATTGATAAACAGTTAGC | Sequence verification |
| *hlyA* qRT-PCR 5' | GCAATGCGGGAAACAGACTC | qPCR |
| *hlyA* qRT-PCR 3' | TTTCTCTGCTGTGCCGAATA | qPCR |
| *cnf1* qRT-PCR 5' | GCTGCTAAGTACCTCCTGGTT | qPCR |
| *cnf1* qRT-PCR 3' | GGTATCTGTTCCGCTTGGACTG | qPCR |
| *hlyA*$^{++}$-*Cm*$^R$ 5' | GCCATATCAATACTGGTGTAGTAACCATCGACGCAACGAAAACTGACGTAGTGTAGGCTGGAGCTGCTTC | Genetic overexpression |
| *hlyA*$^{++}$-*Cm*$^R$ *3'* | ATGGTGTTATTGGTTCCGCGTGCATTGTTATCTATGATACGGGGTCACTAATGGGAATTAGCCATGGTCC | Genetic overexpression |
| *hlyA*$^{++}$ Seq 5' | CCTCTATAACCTGAACATGCTTGG | Sequence verification |
| *hlyA*$^{++}$ Seq 3' | GATGGCGCTGCTCTTCTATTG | Sequence verification |
| *mAPOB* SacI 5' | GATCGAGCTCATGCATTTCCTTTCTTCCTCTTC | Recombinant expression |
| *mAPOB* XhoI 3' | GATCCTCGAGTTCTCCTACTTCAACGTTC | Recombinant expression |
| *Cltc 5'* | CCTTTCTCGCTACTTAGTCCGTC | qPCR |
| *Cltc 3'* | GGTCCTGAGTTTCAGACAAGGC | qPCR |
| *Ap2m1 5'* | CCACAGAACTCAGAGACAGGTG | qPCR |
| *Ap2m1 3'* | CGGCGATACTTGATGCCTTCTC | qPCR |
